# Neurostructural Heterogeneity in Youth with Internalizing Symptoms

**DOI:** 10.1101/614438

**Authors:** Antonia N. Kaczkurkin, Aristeidis Sotiras, Erica B. Baller, Monica E. Calkins, Ganesh B. Chand, Zaixu Cui, Guray Erus, Yong Fan, Raquel E. Gur, Ruben C. Gur, Tyler M. Moore, David R. Roalf, Adon F.G. Rosen, Kosha Ruparel, Russell T. Shinohara, Erdem Varol, Daniel H. Wolf, Christos Davatzikos, Theodore D. Satterthwaite

**Affiliations:** Department of Psychiatry, Perelman School of Medicine, University of Pennsylvania, Philadelphia, PA 19104, USA; Department of Radiology, Perelman School of Medicine, University of Pennsylvania, Philadelphia, PA 19104, USA; Department of Radiology, Washington University, St. Louis, MO, 63130, USA; Center for Biomedical Image Computing and Analytics, University of Pennsylvania, Philadelphia, PA 19104, USA; Philadelphia Veterans Administration Medical Center, Philadelphia, PA 19104; Department of Biostatistics, Epidemiology, and Informatics, University of Pennsylvania, Philadelphia, PA 19104, USA; Department of Statistics, Center for Theoretical Neuroscience, Grossman Center for the Statistics of Mind, Columbia University, New York, NY 10027

**Author notes:** Corresponding author: Antonia N. Kaczkurkin, Ph.D., Richards Building, 5th Floor, Suite 5A, 3700 Hamilton Walk, Philadelphia, PA 19104-6085.

## Abstract

Internalizing disorders such as anxiety and depression are the most common psychiatric disorders, frequently begin in youth, and exhibit marked heterogeneity in treatment response and clinical course. It is increasingly recognized that symptom-based classification approaches to internalizing disorders do not align with underlying neurobiology. An alternative to classifying psychopathology based on clinical symptoms is to identify neurobiologically-informed subtypes based on brain imaging data. We used a recently developed semi-supervised machine learning method (HYDRA) to delineate patterns of neurobiological heterogeneity within youth with internalizing symptoms using structural imaging data collected at 3T from a large community-based sample of 1,141 youth. Using volume and cortical thickness, cross-validation methods indicated a highly stable solution (ARI=.66; permutation-based *p_fdr_* < .001) and identified two subtypes of internalizing youth. Subtype 1, defined by smaller brain volumes and reduced cortical thickness, was marked by impaired cognitive performance and higher levels of psychopathology than both Subtype 2 and typically developing youth. Using resting-state fMRI and diffusion images not considered during clustering, we found that Subtype 1 also showed reduced amplitudes of low-frequency fluctuations in fronto-limbic regions at rest, as well as reduced fractional anisotropy in white matter tracts such as the parahippocampal cingulum bundle and the uncinate fasciculus. In contrast, Subtype 2 showed intact cognitive performance, greater volume, cortical thickness, and amplitudes during rest compared to Subtype 1 and typically developing youth, despite still showing clinically significant levels of psychopathology. Identification of biologically-grounded subtypes of internalizing disorders may assist in targeting early interventions and assessing longitudinal prognosis.

## INTRODUCTION

Internalizing disorders, including depression and anxiety disorders, are the most common psychiatric conditions,^1^ and together result in an enormous worldwide burden of illness.^2^ In contrast to cardiovascular or neurodegenerative disorders, internalizing disorders often begin in youth, leading to a lifetime of morbidity.^1,3^ At present, diagnosis of internalizing disorders remains driven by presenting symptoms, as codified in the *Diagnostic and Statistical Manual of Mental Health Disorders* (*DSM-5*).^4,5^ However, such diagnoses often lack specificity, as evinced by the high degree of comorbidity with other psychiatric disorders^6,7^ and marked heterogeneity in both treatment response and longitudinal outcome.^8^

Emerging evidence from epidemiology, genetics, and clinical neuroimaging often does not support the current diagnoses codified in the DSM.^9–11^ One alternative is to evaluate dimensions of symptoms that cross diagnostic boundaries.^12^ However, dimensional models based on symptoms only do not take into account the neurobiological mechanisms underlying psychiatric symptoms. A classification approach that parses heterogeneous clinical syndromes based on neurobiological data would be a significant advancement for the field.^13^ Intensifying efforts are being made to identify neurobiologically-informed subtypes using machine learning techniques. Using such an approach, patients are clustered into disease sub-groups according to shared patterns in imaging or other data types in order to reveal the heterogeneous biological mechanisms that underlie comorbid disorders. Recent work has delineated neurobiological subtypes in Alzheimer’s disease,^14–16^ depression in adults,^17,18^ and psychosis.^19–21^

To our knowledge, as of yet there have been no efforts to parse neurobiological heterogeneity in youth with internalizing symptoms. Accordingly, the aim of the current study was to delineate patterns of neurobiological heterogeneity in internalizing symptoms in relation to typically developing controls among 1,141 youth using data-driven machine learning techniques. These subtypes were then evaluated using independent clinical, cognitive, and neuroimaging data that was not used in the clustering process.

## METHODS

### Participants

A total of 1,601 participants ages 8-23 years received multi-modal neuroimaging, clinical phenotyping, and cognitive assessment as part of the Philadelphia Neurodevelopmental Cohort (PNC),^22,23^ a large community-based sample of youth. After standard exclusion criteria (including medical disorders and structural image quality; see Supplement), 715 participants met screening criteria for an anxiety and/or depressive disorder and 426 were typically developing youth with no psychiatric diagnoses (n=1,141 total). As expected, the internalizing group showed a greater percentage of females than typically developing youth. Furthermore, modality-specific quality assurance was performed and resulted in a sample of n=840 with resting-state functional MRI (rsfMRI) and n=923 with diffusion imaging data (Supplement).

### Clinical assessment

As described in detail in our previous work^22–24^ and in the Supplement, assessment of lifetime psychopathology was conducted using GOASSESS, a structured screening interview based on a modified version of the K-SADS.^25^ We included participants in the internalizing group if they met criteria for any anxiety and/or depressive disorder including agoraphobia, generalized anxiety disorder, obsessive-compulsive disorder, panic disorder, posttraumatic stress disorder, separation anxiety disorder, social anxiety disorder, specific phobia, or major depressive disorder (**Table 1**).

**Table 1.** Summary of demographic data

|  | **TD**<br>(n=426) |  | **S1**<br>(n=403) |  | **S2**<br>(n=312) |  |
| --- | --- | --- | --- | --- | --- | --- |
|  | *M* | *SD* | *M* | *SD* | *M* | *SD* |
| Age (years) | 14.68 | 4.05 | 15.35 | 3.35 | 15.10 | 3.66 |
|  | N | Percent | N | Percent | N | Percent |
| Gender |  |  |  |  |  |  |
| Female | 208 | 49% | 240 | 60% | 179 | 57% |
| Male | 218 | 51% | 163 | 40% | 133 | 43% |
| Internalizing Disorders (N=715) |  |  |  |  |  |  |
| Agoraphobia | - | - | 51 | 4% | 27 | 2% |
| Generalized Anxiety Disorder | - | - | 13 | 1% | 14 | 1% |
| Major Depression | - | - | 109 | 10% | 82 | 7% |
| Obsessive-Compulsive Disorder | - | - | 26 | 2% | 17 | 1% |
| Panic | - | - | 10 | .9% | 4 | .4% |
| PTSD | - | - | 112 | 10% | 56 | 5% |
| Separation Anxiety | - | - | 29 | 3% | 34 | 3% |
| Social Anxiety | - | - | 193 | 17% | 125 | 11% |
| Specific Phobia | - | - | 230 | 20% | 184 | 16% |
| Co-morbid Disorders |  |  |  |  |  |  |
| ADHD | - | - | 77 | 7% | 70 | 6% |
| Anorexia | - | - | 10 | .8% | 5 | .4% |
| Bulimia | - | - | 2 | .2% | 3 | .3% |
| Conduct Disorder | - | - | 65 | 6% | 19 | 2% |
| Mania | - | - | 7 | .6% | 5 | .4% |
| Oppositional Defiant Disorder | - | - | 200 | 18% | 106 | 9% |
| Psychosis-spectrum | - | - | 169 | 15% | 106 | 9% |

Note. \*Due to comorbidity, individual participants may be present in more than one category of lifetime prevalence.

### Clinical and cognitive factor analyses

As prior,^26,27^ to provide a dimensional summary of the diverse psychopathology data, we used a confirmatory bifactor analysis^28,29^ to model four orthogonal factors (anxious-misery, psychosis, externalizing, and fear) plus a general factor, overall psychopathology, which represents the symptoms common across all psychiatric disorders (Supplement). Cognition was assessed using the University of Pennsylvania Computerized Neurocognitive Battery (CNB).^30^ Fourteen cognitive tests evaluating aspects of cognition were summarized with exploratory factor analysis into three domains: 1) executive function and complex reasoning, 2) social cognition, and 3) episodic memory (Supplement).^30^ Reading skills were measured with the *Wide Range Achievement Test, 4^th^ Edition* (WRAT-4) reading subscale.^31^

### Image acquisition, quality assurance, and image processing

Image acquisition, processing, and quality assurance procedures for volume, cortical thickness, rsfMRI, and DTI measures have been previously described^23,32,33^ and are detailed in the Supplement. Structural images were processed using the top-performing tools included in ANTs.^34^ Functional connectivity among brain regions is primarily attributable to correlations between low-frequency fluctuations in regional activation patterns.^35^ Therefore, we computed the voxel-wise amplitude of low-frequency fluctuations (ALFF)^35^ as the sum over frequency bins in the low-frequency (0.01-0.08 Hertz) band of the power spectrum. Cortical thickness, volume, and ALFF were summarized in anatomic regions within gray matter defined on an individual basis using a top-performing multi-atlas labeling procedure with joint label fusion.^36^ Fractional anisotropy maps were calculated from DTI using FSL^37^ and summarized in tracts defined by the JHU white-matter tractography atlas.^38^

### Parsing heterogeneity with semi-supervised machine learning

To identify neurostructural subtypes within youth with internalizing symptoms, we used a recently developed, semi-supervised machine learning tool, HYDRA (Heterogeneity through Discriminative Analysis).^14^ HYDRA compares cases (internalizing youth) and controls to identify *k* clusters within internalizing youth. In contrast to fully-supervised learning techniques (i.e., support vector machines (SVM) or random forests) which cannot distinguish between subtypes of cases (**Figure 1A**), HYDRA simultaneously performs classification and clustering (**Figure 1B**). Furthermore, in contrast to unsupervised clustering techniques (i.e., k-means or community detection), HYDRA does not cluster patients based on their similarity, which is a process that is vulnerable to confounding inter-individual variations that are irrelevant to disease (e.g., due to age or sex). Instead, HYDRA clusters cases based on their differences from controls. This is accomplished by finding multiple linear hyperplanes, which together form a convex polytope (**Figure 1B**). Rather than coercing participant data points into a single common discriminative pattern, HYDRA allows groups to form based on multiple decision boundaries, thereby rendering the resulting subtypes easily interpretable, compared to other nonlinear classifiers. The result is a data-driven approach to classifying subtypes within internalizing youth that can be evaluated further based on clinical, cognitive, and imaging characteristics.

**Figure 1.**
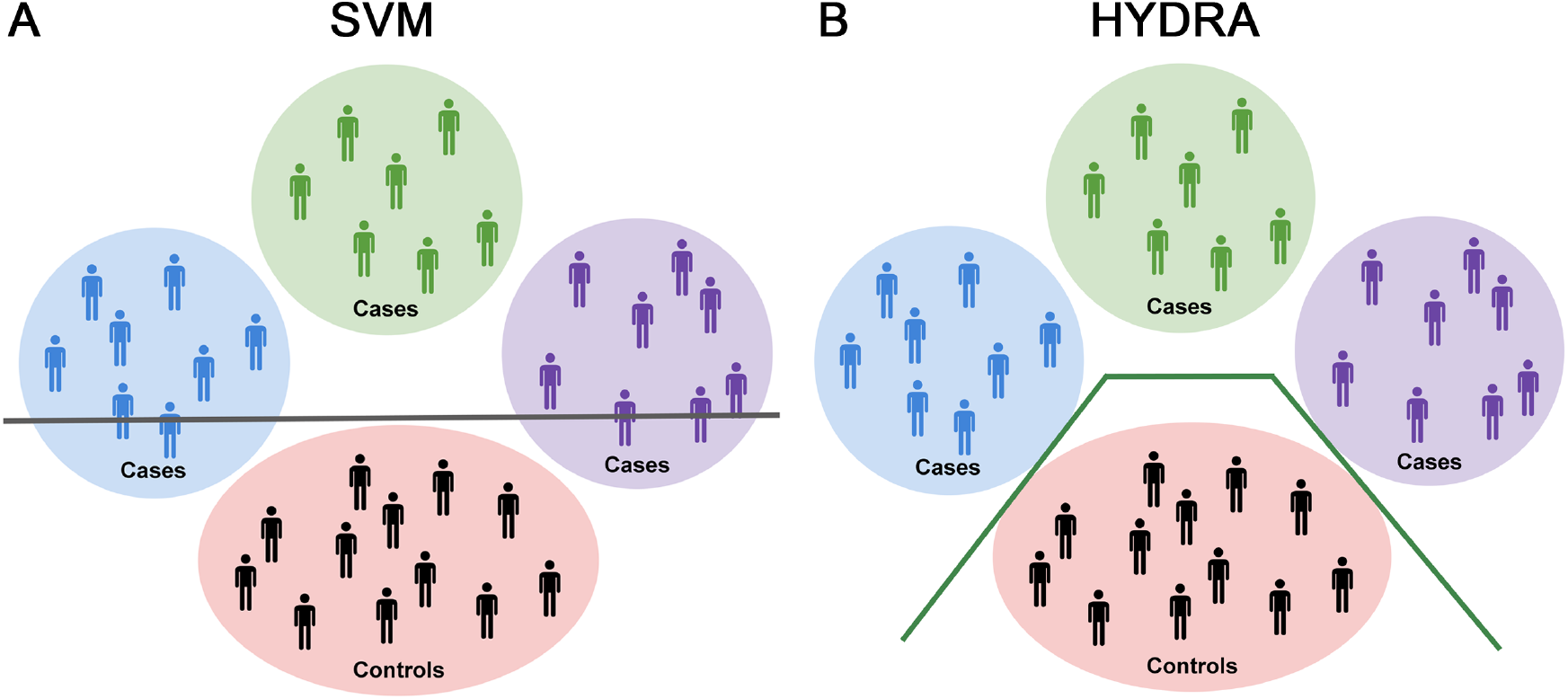
Schematic representing the utility of HYDRA over SVMs for parsing heterogeneity. **A)** Schematic illustrating the use of a linear support vector machine (SVM) to separate cases from controls with a separating hyperplane, shown here as a gray line. Heterogeneity within the cases is represented by the blue, green, and purple circles. As can be seen in this schematic, linear SVMs do not capture the heterogeneity that exists in the cases. **B)** Conversely, HYDRA is able to classify each cluster of cases separately from the controls. This is accomplished by using multiple classifiers that form linear hyperplanes (green lines) whose segments separate the clusters of cases from the controls. The goal is to estimate *k* hyperplanes that distinguish the controls and cases with the largest margin, thus allowing HYDRA to identify heterogeneous groups within the cases.

HYDRA was used to define neurostructural subtypes using the volume of 112 cortical and subcortical regions as well as the cortical thickness of 98 regions, for a total of 210 input features, adjusted for age and sex. Controlling for total brain volume and average cortical thickness before clustering with HYDRA produces clusters with a very low adjusted Rand index (ARI < .20), a measure of out-of-sample reproducibility, suggesting global effects for volume and cortical thickness. Consistent with studies using this technique,^14^ we derived multiple clustering solutions requesting 2 to 10 clusters in order to obtain a range of possible solutions. The ARI was calculated using 10-fold cross validation to evaluate the stability of each solution; the solution with the highest ARI value was selected for subsequent analyses. If instead a one cluster solution exists, then the reproducibility of the solutions will be poor. Permutation testing was used to statistically evaluate the stability of observed ARI values in comparison to a null distribution (see Supplement).

### Group-level statistical analyses

After parsing subtypes of internalizing youth based on structural data, we sought to 1) define how the subtypes differed on psychopathology and cognition, 2) understand what structural features (thickness, volume) drove the subtypes discovered, and 3) investigate differences between the subtypes in two independent neuroimaging sequences not used in clustering (ALFF from rsfMRI and fractional anisotropy from DTI). Both linear and nonlinear age effects were modeled using penalized splines within a generalized additive model, which assesses a penalty on nonlinearity using restricted maximum likelihood (REML) in order to avoid over-fitting.^42,43^ Age, sex, and image quality (see Supplement for details) were modeled as follows:

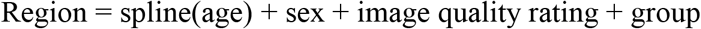

Omnibus ANOVAs and pairwise post-hoc tests were corrected for multiple comparisons by controlling the False Discovery Rate (FDR, *Q*<0.05). Interactions between group and age as well as group and sex were also evaluated. Finally, sensitivity analyses excluding participants who were taking psychotropic medications at the time of imaging (included: n=1,037) were conducted to ensure that our results were not driven by medication effects.

### Data and code availability

See https://github.com/PennBBL/KaczkurkinHeterogenInternalizing for an overview and all data analysis code used in this manuscript. Data from the Philadelphia Neurodevelopmental Cohort can be accessed at https://www.ncbi.nlm.nih.gov/projects/gap/cgi-bin/study.cgi?study_id=phs000607.v3.p2. The HYDRA code can be found at https://github.com/evarol/HYDRA.

## RESULTS

### HYDRA identifies subtypes of internalizing youth with a high degree of stability

HYDRA identified *k* neurostructural subtypes from 210 regional brain features (volume and cortical thickness) after adjusting for age and sex. Evaluation of cluster stability using 10-fold cross-validation exhibited a well-defined peak at *k* = 2 (**Figure 2**), suggesting the existence of two highly reproducible subtypes (ARI = .66) within internalizing youth. Finding a reproducible solution for *k* > 1 suggests that there is structure in the data (in other words, the data is not homogeneous), since the reproducibility of the solution would be poor if the data were instead characterized by a 1 cluster solution. Permutation results further demonstrated a significantly higher ARI for the 2-subtype solution compared to a null distribution (*p_fdr_* < .001).

**Figure 2.**
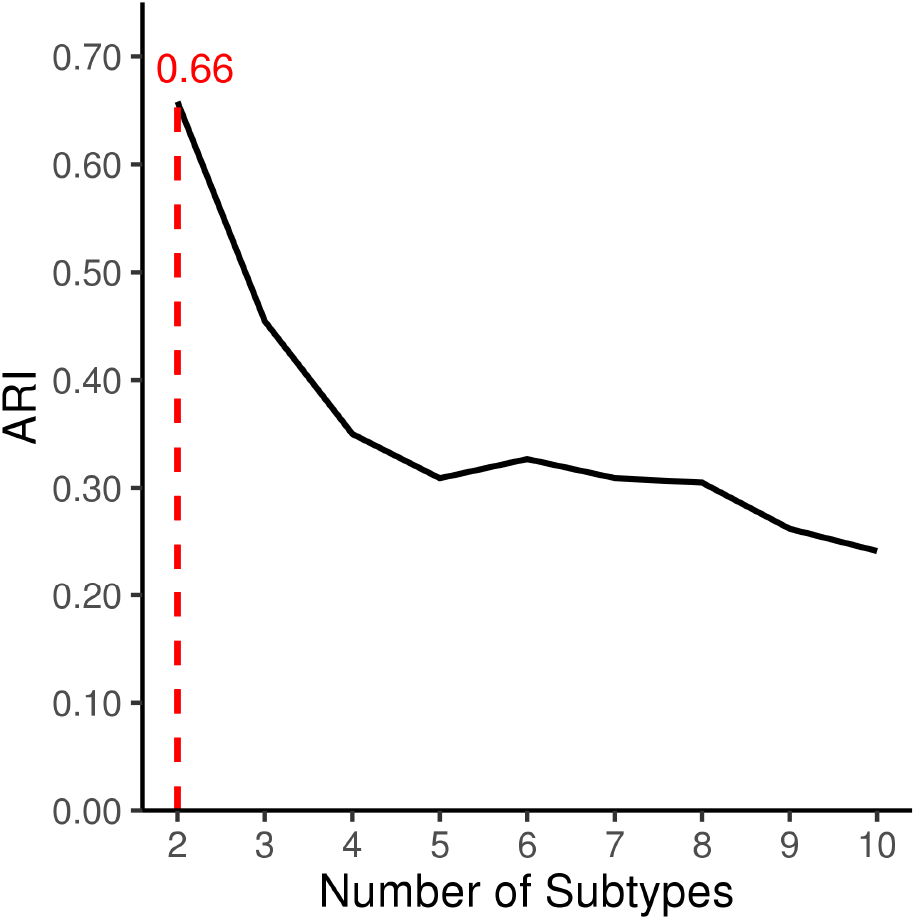
HYDRA identifies 2 subtypes of internalizing youth with a high level of stability. Cross-validated adjusted Rand index (ARI) for 2-10 cluster solutions obtained with HYDRA. The ARI was computed with 10-fold cross-validation to quantify the similarity between different clustering results while controlling for grouping by chance, resulting in a more conservative estimation of the overlap between clustering solutions. The figure shows a clear peak at the 2-cluster solution (shown with a dotted line), suggesting our data has 2 subtypes of internalizing youth which show a high degree of stability (ARI = .66).

### Subtype demographics

As an initial step, we evaluated the demographics of our neurostructural subtypes (**Table S1**). Groups differed in age, with Subtype 1 being slightly older than typically developing youth; no other age effects were significant. All subsequent analyses controlled for age and sex. Subtype 1 also had lower maternal education than Subtype 2 or typically developing youth, while Subtype 2 did not differ from typically developing youth in this regard.

### Elevated psychopathology is present in Subtype 1

Next, we evaluated whether the internalizing neurostructural subtypes differed in terms of psychopathology. Psychopathology symptoms were summarized as factors which reflect anxious-misery, psychosis, behavioral, fear, and overall psychopathology. As expected based on the inclusion criteria (patients all met criteria for an internalizing disorder), Subtype 1 and 2 both showed elevated psychopathology symptoms compared to typically developing youth in all domains (**Table S1**). Accordingly, we focused on differences between the subtypes. Subtype 1 showed greater overall psychopathology symptoms (**Figure 3A**), behavioral symptoms (**Figure 3B**), and fear symptoms (**Figure 3C**) compared to Subtype 2. No significant differences were found between the subtypes for anxious-misery or psychosis. These results demonstrate that Subtype 1 shows a higher burden of psychiatric symptoms than both Subtype 2 and typically developing youth across multiple domains.

**Figure 3.**
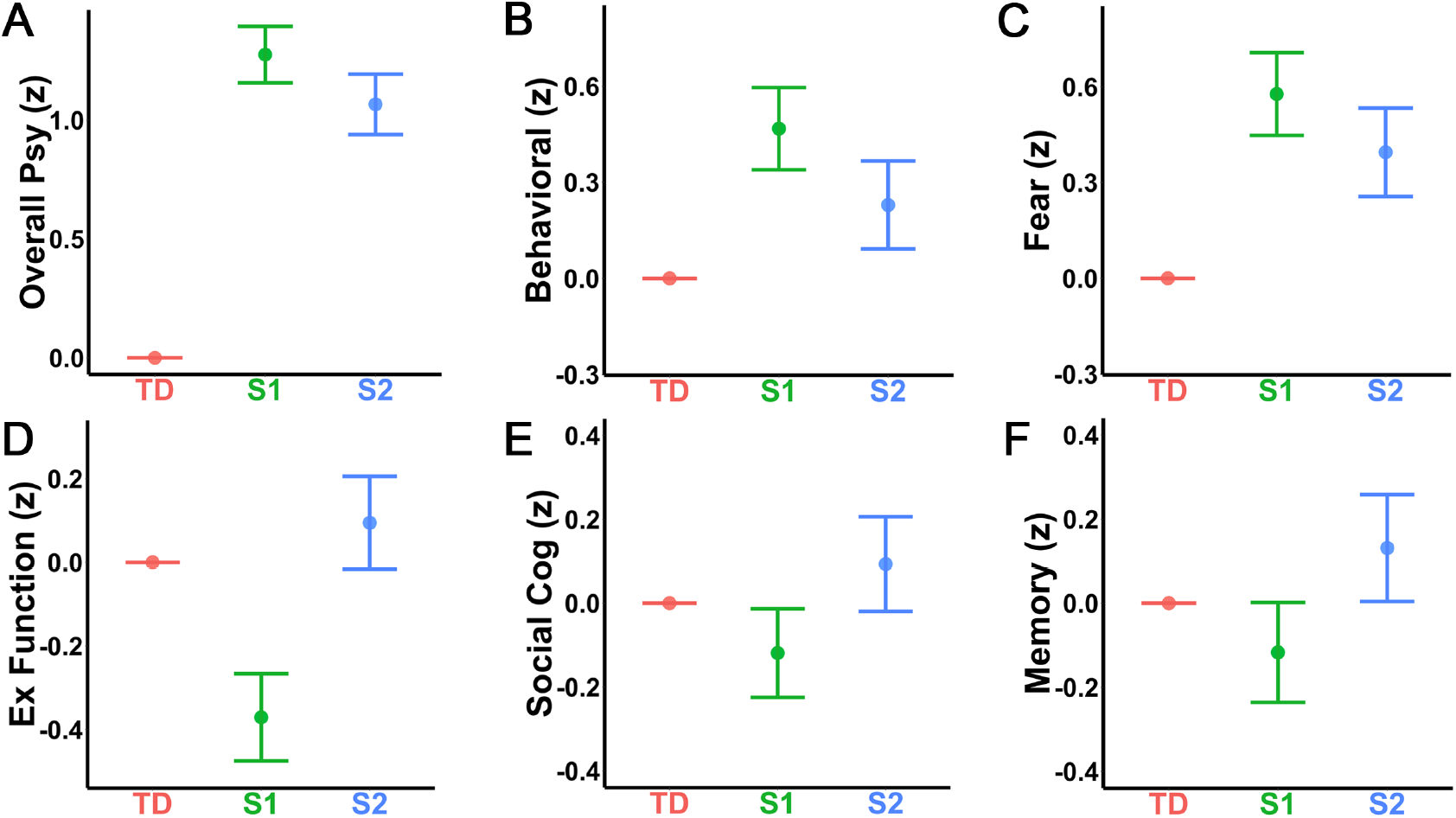
Subtype 1 shows greater psychopathology and poorer cognitive performance. Estimates are shown on the Y-axis from the fitted model testing for group differences in each domain. Each vertical line represents the 95% confidence interval (CI), with the comparison group (typically developing youth: TD) represented by its mean line. The subtype is significantly different from TD if its corresponding CI does not contain 0 (the mean of TD). To examine differences in psychopathology, symptoms were summarized into anxious-misery, psychosis, behavioral, fear, and overall psychopathology factors. As expected, both Subtype 1 (S1) and Subtype 2 (S2) showed greater levels of psychopathology compared to TD across all psychopathology factors, thus, we focus on the differences between S1 and S2. S1 showed higher levels of **A)** overall psychopathology, **B)** behavioral symptoms, and **C)** fear symptoms than S2. There were no significant differences between S1 and S2 for the anxious-misery or psychosis factors. In terms of cognition, S1 showed significantly lower performance on **D)** executive functioning tasks than both TD and S2; S2 did not differ from TD. S1 also performed more poorly relative to the other two groups in terms of **E)** social cognition, while S2 again did not differ from TD. Additionally, S1 performed significantly below S2 on **F)** episodic memory; however, neither S1 nor S2 significantly differed from TD.

### Subtype 1 is marked by impaired cognition

We then evaluated whether the subtypes differed in cognitive performance and reading skills. Notably, this independent data was not used in clustering. Subtype 1 showed significantly reduced overall accuracy relative to Subtype 2 and typically developing youth, with no difference found between typically developing youth and Subtype 2 (**Table S1**). Analyses of the specific accuracy factors revealed that Subtype 1 showed reduced performance relative to Subtype 2 on all factors including executive function/complex reasoning (**Figure 3D**), social cognition (**Figure 3E**), and episodic memory (**Figure 3F**; **Table S1**). Compared to typically developing youth, Subtype 1 had lower performance only for executive function/complex reasoning and social cognition, but not for episodic memory. Subtype 2 did not significantly differ from typically developing youth on any of these measures. In terms of academic skills, Subtype 1 demonstrated lower reading skills than both Subtype 2 and typically developing youth, who did not differ from each other (**Table S1**).

### Subtypes display markedly divergent patterns of brain structure

Having identified two subtypes of internalizing youth based on volume and cortical thickness data, we examined the structural features that drove this clustering. The results demonstrated that Subtype 1 showed smaller regional volumes than either Subtype 2 or typically developing youth for all 112 regions (**Figure 4A** and **S1A**; **Table S2**). Likewise, Subtype 1 had thinner cortex in 96 out of 98 regions compared to Subtype 2 (**Figure 4B; Table S2**) and in 95 regions compared to typically developing youth (**Figure S1B; Table S2**). Compared to typically developing youth, Subtype 2 demonstrated greater volume in 111 regions (**Figure S2A; Table S2**) and greater cortical thickness in 81 regions (**Figure S2B**; **Table S2**). Interactions between group and age as well as group and sex were evaluated but found to be nonsignificant except for volume of occipital fusiform gyrus.

**Figure 4.**
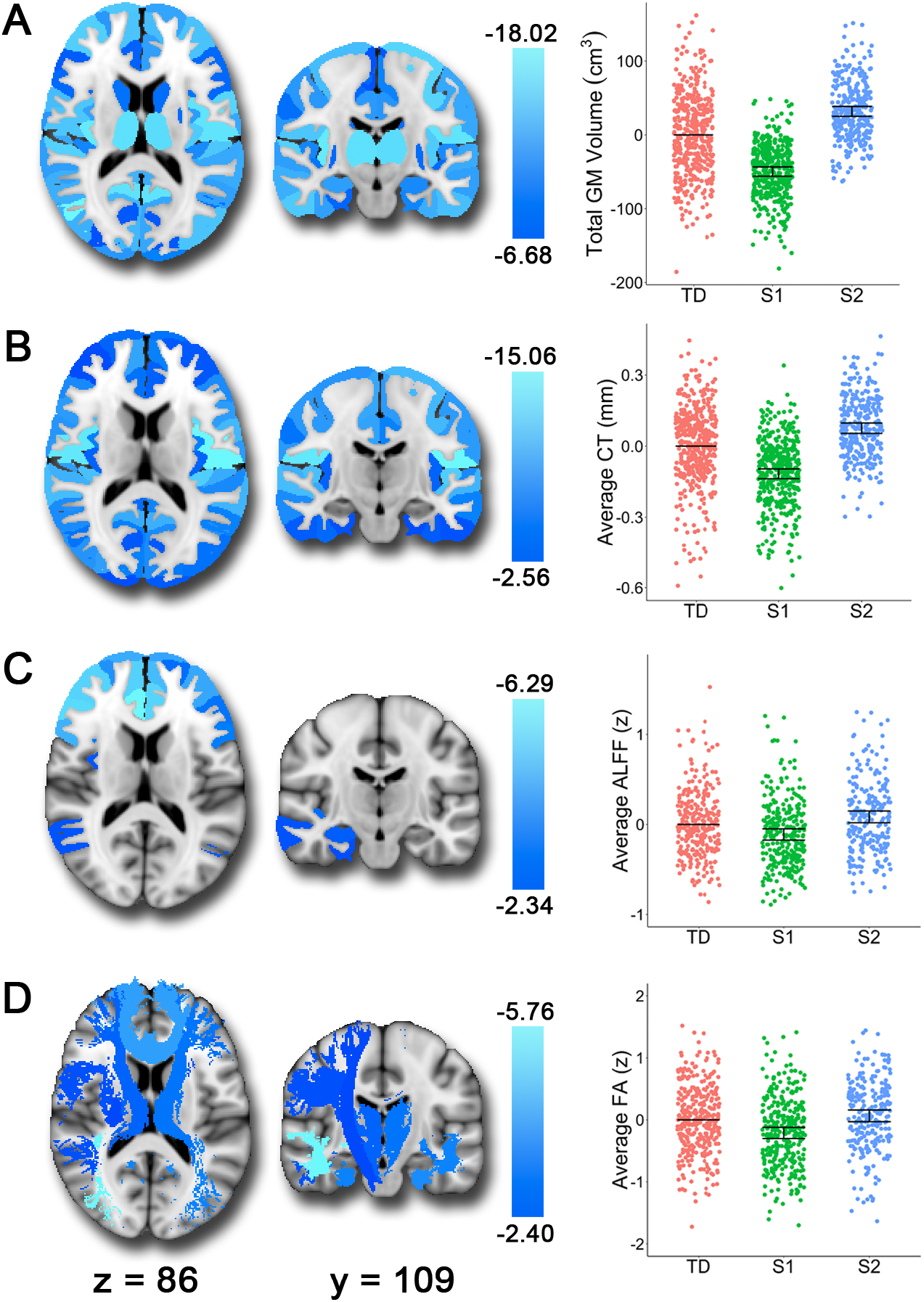
Subtype 1 (S1) shows smaller volume, thinner cortex, lower resting-state ALFF, and reduced white matter integrity relative to Subtype 2 (S2). The brain images show the t-values for the S1>S2 contrast. In the scatterplots, we show the estimates from the fitted GAM model with all three groups for comparison. Each vertical line represents the 95% confidence interval (CI), with the comparison group (typically developing youth: TD) represented by its mean line. The subtype is significantly different from TD if its corresponding CI does not contain 0 (the mean of TD). **A)** S1 showed smaller volumes than S2 consistently across the brain. **B)** In terms of cortical thickness, S1 shows reduced cortical thickness compared to S2 in all regions except the left and right entorhinal cortices. **C)** S1 also demonstrated reduced resting-state ALFF (amplitude of low-frequency fluctuations) in frontal regions, the right amygdala, and the right hippocampus compared to S2. **D)** Finally, relative to S2, we found that S1 showed reduced fractional anisotropy in white matter tracts including the inferior longitudinal fasciculi, uncinate fasciculus, anterior thalamic radiation, corticospinal tract, parahippocampal cingulum bundle, superior longitudinal fasciculus, and forceps minor.

### Subtype 1 shows abnormalities in resting-state and diffusion measures

Finally, to further understand differences in these neurostructural subtypes, we examined two independent imaging modalities which were not used in clustering: ALFF from rsfMRI and fractional anisotropy from DTI. ALFF was reduced in Subtype 1 compared to Subtype 2 in 40 frontal cortex and limbic regions, including bilateral middle/superior frontal gyrus, right amygdala, and right hippocampus (**Figure 4C**; **Table S3**). Subtype 1 also showed reduced amplitudes in 25 of these regions compared to typically developing youth (**Figure S1C**; **Table S3**). Conversely, Subtype 2 demonstrated greater ALFF in 13 regions compared to typically developing youth (**Figure S2C**; **Table S3**). These results suggest that Subtype 1 shows abnormalities in the resting-state power spectrum in regions associated with executive functioning and affective processing.

Differences between Subtype 1 and Subtype 2 were also apparent in fractional anisotropy. Subtype 1 showed reduced fractional anisotropy compared to Subtype 2 in 10 out of 18 white matter tracts including the inferior longitudinal fasciculus, uncinate fasciculus, anterior thalamic radiation, corticospinal tracts, parahippocampal cingulum bundle, superior longitudinal fasciculi, and forceps minor (**Figure 4D; Table S4**). Subtype 1 also showed reduced fractional anisotropy in 8 tracts compared to typically developing youth (**Figure S1D; Table S4**). Subtype 2 showed relatively similar fractional anisotropy to typically developing youth, with only the left and right anterior thalamic radiations demonstrating higher levels in Subtype 2 (**Figure S2D; Table S4**). These results emphasize that Subtype 1 has reduced white matter integrity in several key tracts that link frontal cortex and limbic brain regions.

### Sensitivity analyses provide convergent results

Sensitivity analyses were conducted after excluding the minority of participants who were taking psychotropic medications at the time of imaging (included: n=1,037). In this subsample, the pattern of results for demographics, psychopathology, and cognition/academic skills remained highly similar (**Table S5**). For the structural results, sensitivity analyses yielded nearly identical results (**Table S6**). In addition, 27 out of 41 resting-state ALFF regions remained significant between the subtypes (**Table S7**). The fractional anisotropy results were quite similar, with 9 out of 10 tracts remaining significant between Subtype 1 and 2 (**Table S8**).

## DISCUSSION

Capitalizing on a large sample of youth and recent advances in semi-supervised machine learning, we identified two reliable neurostructural subtypes of internalizing disorders. Subtype 1 was marked by elevated levels of psychopathology, impaired cognition, and multiple deficits apparent on multi-modal imaging. These deficits included distributed loss of gray matter volume and cortical thickness, reduced ALFF in fronto-limbic cortex, and reduced integrity of white matter tracts. In contrast, Subtype 2 had preserved cognitive functioning and brain integrity despite clinically significant levels of psychopathology. These results provide a new account of the heterogeneity in brain structure and function present in youth with internalizing disorders.

### Heterogeneous neurostructural abnormalities in internalizing disorders

The pattern of deficits revealed between the subtypes of internalizing youth illustrates the detrimental effects associated with abnormal structural development. Reduced volume and cortical thickness have been associated with deficits in cognitive functioning,^46,47^ impaired academic skills,^47–49^ and greater psychopathology.^50–55^ These structural abnormalities are likely the result of a combination of genetic and environmental effects.^56^ Environmental factors, such as low SES and childhood adversity, are associated with chronic exposure to stress hormones,^57,58^ which have been shown to impact the development of structures related to psychopathology^59,60^ and cognition.^61,62^ In line with this prior research, we found an association between neurostructural deficits and both impaired cognition and greater levels of psychopathology. This is consistent with prior work from our group and others showing a robust relationship between psychopathology and structural brain deficits.^50–55^

We expand on previous work by showing related deficits in two independent modalities, with neurostructural deficits associated with reduced resting-state ALFF and fractional anisotropy in white matter tracts. Reduced resting-state ALFF may reflect dysregulated regional spontaneous neural activity.^35^ The fronto-limbic pattern of reduced ALFF observed in the current study suggests potential deficits in executive functioning regions, which is in line with the poorer cognitive performance we found in those with reduced volume and cortical thickness. Our results are also consistent with prior studies showing reduced resting-state ALFF in frontal regions in children with ADHD.^35^ Additionally, the finding of reduced white matter integrity in tracts such as the inferior longitudinal fasciculus, uncinate fasciculus, and forceps minor is consistent with previous studies showing a relationship between decreased fractional anisotropy in these tracts and greater depressive^63,64^ and ADHD symptoms.^65,66^ Similarly, reduced fractional anisotropy has also been linked to poorer cognitive functioning.^67^

In contrast to the deficits seen in Subtype 1, Subtype 2 was characterized by preserved brain structure and cognitive functioning, but still showed high levels of psychopathology. This may suggest compensatory mechanisms, whereby individuals with greater brain reserve can compensate for deficits typically associated with psychopathology, allowing for preserved cognition.^68^ Together, these results demonstrate the impact of abnormal structural development on cognitive and affective functioning, with greater neural resources potentially mitigating detrimental effects on cognition. However, this does not explain why preserved brain structure, function, and cognition does not also protect against psychiatric symptoms, suggesting disparate pathways to apparently similar manifestations of psychopathology.^69^

### Advances in parsing neurobiological heterogeneity in youth

The results of the current study provide both conceptual and methodological advances in the classification of internalizing disorders in youth. Prior studies have primarily used symptom-based diagnostic categories to explore associated neurobiological mechanisms in a case-control design, or examined associations between dimensional clinical phenotypes and imaging measures across diagnostic categories. However, several recent efforts have sought to define subtypes in psychiatric disorders using clustering techniques. For example, our collaborative group has previously used machine learning techniques to identify subtypes of Alzheimer’s disease characterized by distinct atrophy patterns in the brain.^14–16^ Similar approaches have been used to parse heterogeneity in adults with depression using resting-state connectivity data.^17,18^ Likewise, subtypes of psychosis have been identified using multimodal data;^19–21^ however, two of these studies derived their subtypes using psychiatric symptoms rather than brain measures.^20,21^ While clustering methods have been used previously to identify subtypes of internalizing adults,^70^ these methods have clustered on symptoms only and then related symptom subtypes to neuroimaging measures. The current study extends this research by using structural data as input for clustering to delineate neuroanatomical subtypes of internalizing symptoms, which were then evaluated using independent cognitive and neuroimaging measures. Additionally, while the majority of prior studies on heterogeneity have considered adults, our results build upon this work by parsing neurostructural heterogeneity in a community-based sample of youth.

In addition to studying youth, this study is distinctive from prior efforts in its methodological approach. Until relatively recently, the majority of approaches for understanding the neurobiological differences underlying psychiatric symptoms (case-control, linear SVMs) have assumed that a single discriminative pattern differentiates those with psychiatric symptoms from healthy controls. However, as noted above, there has been a movement towards using methods that parse neurobiological heterogeneity within patient groups.^14–21^ While unsupervised methods cluster patients based on how similar they are to each other, one advantage of the approach taken here is that the semi-supervised learning procedure implemented in HYDRA allows us to cluster patients by how different they are from controls, yielding subtype-specific neurobiological signatures that differ from controls. In contrast, traditional clustering techniques are susceptible to splitting the data on non-specific factors such as age or sex, producing clusters that may not be aligned well on psychopathology. In the current study, HYDRA allowed us to identify two subtypes of youth with internalizing disorders that differed from controls on multiple clinical, cognitive, and imaging measures of interest.

## CONCLUSIONS

This study provides important data regarding the neurostructural heterogeneity in youth with internalizing disorders. We identified two subtypes of internalizing youth differentiated by abnormalities in brain structure, function, and white matter integrity, with one subtype showing poorer functioning across multiple domains. Given the early age of onset found in anxiety and depressive symptoms during development,^3^ biomarkers that dissect heterogeneous neural patterns in a developmental sample may aid in identifying youth at risk for these symptoms. A greater comprehension of how abnormalities in the brain give rise to these symptoms in youth is critical for the development of earlier and more effective treatments that may reduce the negative long-term outcomes associated with internalizing symptoms.

## ACKNOWLEDGEMENTS

This work was supported by grants from the National Institute of Mental Health (RC2 grants MH089983 and MH089924 to REG, K99MH117274 to ANK, R01MH107703 and R01MH113550 to TDS, R01NS085211 to RTS, R01MH112847 to RTS and TDS, R01MH107235 to RCG, R01MH11365 to DHW, and R01MH112070 and R01EB022573 to CD), the Dowshen Program for Neuroscience, the Lifespan Brain Institute at the Children’s Hospital of Philadelphia and Penn Medicine, the NARSAD Young Investigator Award (ANK), and the Center for Biomedical Computing and Image Analysis (CBICA) at Penn for developing statistical analyses (RTS & TDS) and multivariate pattern analysis software (AS & TDS).

