## Supplement for "Neurostructural Heterogeneity in Youth with Internalizing Symptoms"

### SUPPLEMENTARY MATERIALS

#### Participants

A total of 1,601 participants 8-23 years of age received multi-modal neuroimaging, clinical phenotyping, and cognitive assessment as part of the Philadelphia Neurodevelopmental Cohort (PNC),[1,2] a large community-based sample of youth. From the pool of 1,601 participants, 154 were ineligible for the current study due to: 1) medical disorders that could impact brain functioning (n=81), 2) medication use that could affect central nervous system functioning (n=64), 3) incidentally discovered structural brain abnormalities[3] (n=20), or 4) failing to meet structural image quality assurance protocols[4] (n=51); several subjects were ineligible for multiple criteria. Additionally, two participants were missing clinical diagnostic data, 17 were missing data on maternal level of education, and three were missing cognitive data. These participants were excluded from all analyses.

#### Clinical assessment

As described in detail in our previous work,[1,2,5] assessment of lifetime psychopathology was conducted using GOASSESS, a structured screening interview administered to probands (age 11-21) and collateral informants of probands (age 8-17), based on a modified version of the *Kiddie-Schedule for Affective Disorders and Schizophrenia*[6] and *Diagnostic and Statistical Manual of Mental Disorders, 4th edition, Text Revision criteria*.[7] The GOASSESS interview assesses lifetime occurrence of mood (major depressive episode, mania), anxiety (agoraphobia, generalized anxiety, panic, specific phobia, social phobia, separation anxiety, posttraumatic stress), behavioral problems (oppositional defiant, attention deficit/hyperactivity, conduct), psychosis, eating disorder (anorexia, bulimia), and suicidal symptoms. Of note, due to comorbidity, participants may be represented in more than one diagnostic category. The GOASSESS interview was administered by trained assessors who underwent a common training protocol (developed and implemented by Dr. Calkins) that included didactic sessions, assigned readings, and supervised pair-wise practice. Assessors were certified for independent assessments following observation by a certified clinical observer who rated the proficiency of the assessor on a 60-item checklist of interview procedures. The median interval of time between clinical assessment and neuroimaging was 2 months.

#### Clinical and cognitive factor analyses

To provide a dimensional summary of the psychopathology data, we applied an exploratory factor analysis to 112 item-level symptom questions from the GOASSESS interview, which has been described in detail elsewhere.[8] This exploratory factor analysis yielded four correlated dimensions of psychopathology including factors for anxious-misery, psychosis, behavioral (externalizing), and fear. We then used a confirmatory bifactor analysis[9,10] implemented in Mplus[11] to orthogonally model the four factors plus overall psychopathology, which represents the symptoms common across all psychiatric disorders. These orthogonal factors from the bifactor analysis were used in the present study. For the distribution of the individual GOASSESS interview items within the factors, see our previous publication.[8]

Cognition was assessed using the University of Pennsylvania Computerized Neurocognitive Battery (CNB), which has been described previously.[12] Briefly, 14 cognitive tests evaluating aspects of cognition, including executive control, episodic memory, complex reasoning, social cognition, and sensorimotor speed, were administered in a fixed order. Except for two sensorimotor tests that only measure speed, each test provides measures of both accuracy and speed. We used a measure of overall accuracy, which is the general factor derived from a previously reported bifactor analysis and represents the total accuracy across the 12 accuracy scores.[12] To better understand differences in overall accuracy, we also used three factor scores for performance accuracy derived in a previously reported[12] exploratory factor analysis with oblique rotation: 1) executive function and complex reasoning, 2) social cognition, and 3) episodic memory. Reading skills were measured with the *Wide Range Achievement Test, 4th Edition* (WRAT-4) reading subscale with scores reported as T-scores.[13]

**Image acquisition and processing**

*Volume and cortical thickness imaging*

Imaging data were acquired on a Siemens TIM Trio 3 Tesla scanner (Erlangen, Germany) with a 32-channel head coil. Structural brain imaging was completed using a magnetization-prepared, rapid acquisition gradient-echo (MPRAGE) T1-weighted image with the following parameters: TR = 1810 ms; TE = 3.51 ms; FoV = 180 x 240 mm; matrix 192 x 256; 160 slices; slice thickness/gap 1/0 mm; TI 1100 ms; flip angle 9 degrees; effective voxel resolution of 0.93 x 0.93 x 1.00 mm; total acquisition time 3:28 minutes. Structural image processing utilized Advanced Normalization Tools (ANTs).[14] This pipeline includes N4 bias field correction, brain extraction, Atropos probabilistic tissue segmentation,[15] and direct estimation of cortical thickness in volumetric space.[16] Structural images were registered to a custom template using the top-performing SyN diffeomorphic registration method.[17] In order to parcellate the brain into anatomically-defined regions, we used an advanced multi-atlas labelling approach. Specifically, 24 young adult T1 images from the OASIS data set that were manually labeled by Neuromorphometrics, Inc. (http://Neuromorphometrics.com/) were registered to each subject's T1 image using the top-performing SyN diffeomorphic registration.[17,18] These label sets were synthesized into a final subject-level parcellation using the top-performing joint label fusion (JLF) segmentation procedure.[19]

*Resting-state fMRI*

Approximately 6 minutes of task-free functional data were acquired for each subject using a blood oxygen level-dependent (BOLD-weighted) sequence (TR = 3000 ms; TE =32 ms; FoV = 192 × 192 mm; resolution 3 mm isotropic; 124 spatial volumes). Subjects were instructed to stay awake, keep their eyes open, fixate on the displayed crosshair, and remain still. Task-free functional images were processed using previously described methods.[20] Briefly, preprocessing included 1) correction for distortions induced by magnetic field inhomogeneities using FSL's FUGUE utility, 2) removal of the 4 initial volumes of each acquisition, 3) realignment of all volumes to a selected reference volume using MCFLIRT,[21] 4) demeaning and removal of any linear or quadratic trends, 5) co-registration of functional data to the high-resolution structural image using boundary-based registration,[22] and 6) temporal filtering using a first-order Butterworth filter with a passband between 0.01 and 0.08 Hz. These preliminary processing stages were then followed by the confound regression procedures, as described previously.[20] In order to prevent frequency-dependent mismatch during confound regression,[23] all regressors were band-pass filtered to retain the same frequency range as the data.

Functional connectivity among brain regions is primarily attributable to correlations between low-frequency fluctuations in regional activation patterns. The voxel-wise amplitude of low-frequency fluctuations (ALFF)[24] was computed as the sum over frequency bins in the low-frequency (0.01-0.08 Hertz) band of the voxel-wise power spectrum, computed using a Fourier transform of the time-domain of the voxel-wise signal. ALFF was calculated on data smoothed in SUSAN[25] using a Gaussian-weighted kernel with 6mm FWHM.

*Diffusion tensor imaging*

DTI scans were acquired using a twice-refocused spin-echo (TRSE) single-shot echo-planar imaging (EPI) sequence (TR = 8100 ms; TE = 82 ms; FoV = 240 x 240 $mm^2$; Matrix = RL:128/AP:128/Slices:70, in-plane resolution (x and y) 1.875 $mm^2$; slice thickness = 2mm, gap = 0; flip angle = 90º/180º/180º, volumes = 71, GRAPPA factor = 3, bandwidth = 2170 Hz/pixel, PE direction = AP). This sequence used a four-lobed diffusion encoding gradient scheme combined with a 90-180-180 spin-echo sequence designed to minimize eddy-current artifacts. DTI data were acquired in two consecutive series consisting of 32 diffusion encoding gradient schemes. The complete sequence consisted of 64 diffusion weighted directions with b=1000 s/$mm^2$ and 7 interspersed scans where b=0 s/$mm^2$. The duration of DTI scans was approximately 11 minutes. The imaging volume was prescribed in axial orientation covering the entire cerebrum with the topmost slice just superior to the apex of the brain.[2] In addition to the DTI scan, a map of the main magnetic field (i.e., B0) was derived from a double-echo, gradient-recalled echo (GRE) sequence, allowing us to estimate field distortions in each dataset.

Two consecutive 32-direction acquisitions were merged into a single 64-direction timeseries. The skull was removed for each subject by registering a binary mask of a standard fractional anisotropy (FA) map (FMRIB58 FA) to each subject's DTI image using a rigid body transformation. Eddy currents and subject motion were estimated and corrected using FSL's eddy tool.[26] Diffusion gradient vectors were then rotated to adjust for subject motion estimated by eddy. After the field map was estimated, distortion correction was applied to DTI data using FSL's FUGUE.[27] Lastly, the diffusion tensor and fractional anisotropy were estimated at each voxel using the DTIFIT procedure in FSL's Diffusion Toolbox (FDT).[28]

Registration from native space to a template space was completed using DTI-TK.[29,30] First, the DTI outputs (e.g. FA maps) of DTIFIT were converted to DTI-TK format. Next, a template was generated from the tensor volumes using 14 representative diffusion data sets that were considered "excellent" from the PNC sample; the details of this procedure are published.[31] Ultimately, one high-resolution refined template was created and used for registration of the remaining diffusion datasets. All DTI maps were then registered to the high-resolution study-specific template using DTI-TK. Standard regions of interest (ICBM-JHU White Matter Tracts) were registered from MNI152 space to the study-specific template using ANTs registration.[17] Finally, mean FA was calculated over each white matter tract.

**Image quality assurance**

*Volume and cortical thickness QA*

Three highly trained image analysts independently assessed structural image quality; for full details of this procedure see Rosen et al.[32] Briefly, three raters were trained prior to rating images on an independent training sample of 100 images. All three raters were trained to >85% concordance with faculty consensus ratings. T1 images were rated on a 0-2 Likert scale (0 = unusable images, 1 = usable images with some artifact, and 2 = images with none or almost no artifact). All images with an average rating of 0 were excluded from analyses. We included average quality rating across the three raters as a covariate in the models in order to control for the confounding influence of subtle variation in image quality.

All processed data underwent rigorous quality control as well. Specifically, the volume and thickness of anatomically-defined regions of interest (defined using multi-atlas labeling with joint label fusion) were evaluated for outliers. Outliers were defined as values greater or less than 2.5 standard deviations (SD) from the mean regional value. Participants with an elevated (+2.5 SD) number of regions with outlying volume or cortical thickness values were flagged for manual review. Similarly, a regional laterality index was calculated for both cortical thickness and volume, and participants with an elevated number of regional laterality outliers (+2.5 SD) were flagged for review. Flagged images were then manually viewed by two independent data analysts.

*Resting-state fMRI QA*

As part of the processing procedure for resting BOLD imaging data, the quality of all acquired images was assessed according to a number of metrics. For each of five quality metrics, a minimal inclusion threshold was established and subjects that failed to meet this threshold were omitted from the final sample. Reasons for exclusion/metrics of quality included: 1) resting data not acquired, 2) number of frames with motion exceeding 0.25 mm (> 20 frames), and 3) mean relative RMS displacement (> 0.2 mm framewise). Voxel-wise coverage was assessed on a regional level. Subject motion was assessed on the basis of outputs from the MCFLIRT motion realignment procedure.[21] Two criteria for sample inclusion were obtained from the MCFLIRT output: mean relative displacement and the number of motion spikes. The mean relative displacement indicates the average volume-to-volume motion of the subject according to the root-mean-square metric. The number of motion spikes indicates the number of single frames with relative motion in excess of 0.25 mm according to the root-mean-square metric. For the purposes of resting sample inclusion, hard thresholds were established at 0.2 mm for mean relative RMS displacement and 20 frames for number of spikes. In addition to this exclusion criteria, we used a continuous measure of data quality as a covariate in our statistical analyses. As in our prior work,[33,34] the primary summary metric of subject motion used for resting-state data was the mean relative RMS (root-mean-squared) displacement calculated during time series realignment using MCFLIRT. This metric was included as a covariate to all resting-state analyses.

*Diffusion-weighted imaging QA*

Diffusion data quality assurance has been described in detail elsewhere.[31] Briefly, this process involved initial visual inspection which was done to ensure data fidelity. Individuals with structural abnormalities were excluded. Additionally, any scans collected without the default scan parameters were excluded. Data passing this initial quality assurance step then underwent manual inspection by a trained analyst who inspected each diffusion series in detail. This manual evaluation determined whether each image would be included.

As prior,[31] in addition to this manual evaluation, automated measures of image quality were calculated and used as a covariate in statistical testing. Specifically, the temporal signal-to-noise ratio (SNR) was estimated for each brain voxel using only the 64 b=1000 s/mm2 DTI volumes. The voxel-wise SNR was calculated from the mean and standard deviation of each voxel's intensity, after brain masking and motion correction was measured. Subsequently, the average of all brain voxel temporal SNR was calculated to report a single metric of overall scan SNR, which was used as an additional covariate in diffusion analyses.

#### Group-level statistical analyses

After parsing subtypes of internalizing youth based on structural data, we sought to 1) define how the subtypes differed on psychopathology and cognition, 2) understand what structural features (thickness, volume) drove the subtypes discovered, and 3) investigate differences between the subtypes in two independent neuroimaging sequences not used in clustering (ALFF from rsfMRI and fractional anisotropy from DTI). Given that brain development is a non-linear process,[34–36] we modeled both linear and nonlinear age effects using penalized splines within a generalized additive model (GAM), which assesses a penalty on nonlinearity using restricted maximum likelihood (REML) in order to avoid over-fitting.[37,38] Based on known sex differences in structure,[39] we included sex as a covariate in the model. As expected given that internalizing disorders are more common in females, the subtypes showed a greater percentage of females than typically developing youth. Additionally, quality ratings for each imaging modality were added as an additional model covariate to control for the potential confounding effects of image quality (Supplement).[32] Our group variable was modeled as a factor with three levels: typically developing, Subtype 1 and Subtype 2. We examined group differences in each brain region as follows:

Region = spline(age) + sex + image quality rating + group

Omnibus ANOVAs testing for group differences were corrected for multiple comparisons by controlling the False Discovery Rate (FDR, $Q<0.05$). We then conducted pairwise post-hoc tests (S1 vs. TD, S2 vs. TD, and S1 vs. S2) to determine which groups significantly differed from each other, which were also corrected for multiple comparisons using FDR. Interactions between group and age as well as group and sex were also evaluated. Finally, sensitivity analyses excluding participants who were taking psychotropic medications at the time of imaging (excluded: n=104; included: n=1,037) were conducted to ensure that our results were not driven by medication effects.

#### Permutation testing of subtypes

Permutation tests are useful tools for assessing statistical significance when the underlying distribution of data is unknown.[40] Here we define the null distribution of the subtypes using the healthy control sample, where disease-related variability is not present. For this analysis, half of the healthy controls (n = 426) were randomly assigned to the control group (n=213) and half to the pseudo-patient group (n=213), and these samples were permuted 75 times in HYDRA. We then compared these results to a real-patient group of equal number that included the control samples and real-patient samples (each also equal to n=213). The control and real-patient samples were also permuted 75 times in HYDRA. The clustering stability defined by the adjusted Rand index (ARI) was compared between the subtypes derived from the pseudo-patient (null) and real-patient samples. The ARI for the 2-cluster solution was significantly higher in the real-patient sample compared to that of the null distribution (i.e., pseudo-patient; $p_{fdr} < .001$). The ARIs for the other cluster solutions were not different from the null distribution ($p_{fdr} > 0.05$), except for the 7-cluster solution ($p_{fdr} = 0.014$); however, the stability of 7-cluster solution was substantially lower (ARI of .30). Overall, the permutation testing demonstrated that the 2-cluster solution is significantly more stable than that expected by chance alone.

23 Hallquist MN, Hwang K, Luna B. The nuisance of nuisance regression: Spectral misspecification in a

common approach to resting-state fMRI preprocessing reintroduces noise and obscures functional connectivity. *Neuroimage* 2013; **82**: 208–225.

24 Yu-feng Z, Yong H, Chao-zhe Z, Qing-jiu C, Man-qiu S, Meng L *et al.* Altered baseline brain activity in children with ADHD revealed by resting-state functional MRI. *Brain Dev* 2007; **29**: 83–91.

25 Smith SM, Brady JM. SUSAN - A new approach to low level image processing. *Int J Comput Vis* 1997; **23**: 45–78.

26 Andersson JLR, Sotiropoulos SN. An integrated approach to correction for off-resonance effects and subject movement in diffusion MR imaging. *Neuroimage* 2016; **125**: 1063–1078.

27 Jenkinson M, Beckmann CF, Behrens TEJ, Woolrich MW, Smith SM. FSL. *Neuroimage* 2012; **62**: 782–790.

28 Behrens TEJ, Berg HJ, Jbabdi S, Rushworth MFS, Woolrich MW. Probabilistic diffusion tractography with multiple fibre orientations: What can we gain? *Neuroimage* 2007; **34**: 144–155.

29 Zhang H, Yushkevich PA, Alexander DC, Gee JC. Deformable registration of diffusion tensor MR images with explicit orientation optimization. *Med Image Anal* 2006; **10**: 764–785.

30 Zhang B, Xu Y, Zhu B, Kantarci K. The role of diffusion tensor imaging in detecting microstructural changes in prodromal Alzheimer's disease. *CNS Neurosci Ther* 2014; **20**: 3–9.

31 Roalf DR, Quarmley M, Elliott MA, Satterthwaite TD, Vandekar SN, Ruparel K *et al.* The impact of quality assurance assessment on diffusion tensor imaging outcomes in a large-scale population-based cohort. *Neuroimage* 2016; **125**: 903–919.

32 Rosen AFG, Roalf DR, Ruparel K, Blake J, Seelaus K, Villa LP *et al.* Quantitative assessment of structural image quality. *Neuroimage* 2018; **169**: 407–418.

33 Satterthwaite TD, Elliott MA, Gerraty RT, Ruparel K, Loughead J, Calkins ME *et al.* An improved framework for confound regression and filtering for control of motion artifact in the preprocessing of resting-state functional connectivity data. *Neuroimage* 2013; **64**: 240–256.

34 Satterthwaite TD, Shinohara RT, Wolf DH, Hopson RD, Elliott MA, Vandekar SN *et al.* Impact of puberty on the evolution of cerebral perfusion during adolescence. *Proc Natl Acad Sci* 2014; **111**: 8643–8648.

35 Giedd JN, Blumenthal J, Jeffries NO, Castellanos FX, Liu H, Zijdenbos A *et al.* Brain development during childhood and adolescence: a longitudinal MRI study. *Nat Neurosci* 1999; **2**: 861–863.

36 Lenroot RK, Gogtay N, Greenstein DK, Wells EM, Wallace GL, Clasen LS *et al.* Sexual dimorphism of brain developmental trajectories during childhood and adolescence. *Neuroimage* 2007; **36**: 1065–1073.

37 Wood SN. Stable and efficient multiple smoothing parameter estimation for generalized additive models. *J Am Stat Assoc* 2004; **99**: 673–686.

38 Wood SN. mgcv: GAMs and generalized ridge regression for R. *R News* 2001; **1**: 20–25.

39 Kaczkurkin AN, Raznahan A, Satterthwaite TD. Sex differences in the developing brain: insights from multimodal neuroimaging. *Neuropsychopharmacology* 2019; **44**: 71–85.

40 Nichols TE, Holmes AP. Nonparametric Permutation Tests for Functional Neuroimaging: A Primer with Examples. *Hum Brain Mapp* 2001; **15**: 1–25.

**Supplementary Table 1. Group differences on demographics, cognition, academic achievement, and psychopathology (n=1,141).**

| | | | S1-TD | | | | S2-TD | | | | S1-S2 | | | |
|---|---|---|---|---|---|---|---|---|---|---|---|---|---|---|
| | ***F/X²*** | ***p*** | ***B*** | ***SE*** | ***t*** | $p_{fdr}$ | ***B*** | ***SE*** | ***t*** | $p_{fdr}$ | ***B*** | ***SE*** | ***t*** | $p_{fdr}$ |
| Age* | 3.48 | .031 | 0.67 | 0.26 | 2.61 | .027 | 0.42 | 0.28 | 1.51 | .196 | 0.26 | 0.28 | 0.91 | .361 |
| Gender* | 10.64 | .005 | 0.43 | 0.14 | 3.09 | .006 | 0.34 | 0.15 | 2.29 | .033 | 0.09 | 0.15 | 0.59 | .557 |
| Maternal level of education* | 20.56 | <.001 | -1.01 | 0.17 | -5.99 | <.001 | -0.12 | 0.18 | -0.67 | .503 | -0.89 | 0.18 | -4.86 | <.001 |
| Academic achievement^ | 35.56 | <.001 | -8.00 | 1.10 | -7.25 | <.001 | 0.61 | 1.18 | 0.51 | .608 | -8.61 | 1.19 | -7.23 | <.001 |
| Overall accuracy^ | 28.47 | <.001 | -0.29 | 0.05 | -5.68 | <.001 | 0.10 | 0.05 | 1.82 | .069 | -0.38 | 0.05 | -7.07 | <.001 |
| Accuracy factors^ | ***F*** | $p_{fdr}$ | ***B*** | ***SE*** | ***t*** | $p_{fdr}$ | ***B*** | ***SE*** | ***t*** | $p_{fdr}$ | ***B*** | ***SE*** | ***t*** | $p_{fdr}$ |
| Executive function/complex reasoning | 39.26 | <.001 | -0.37 | 0.05 | -6.97 | <.001 | 0.09 | 0.06 | 1.67 | .096 | -0.47 | 0.06 | -8.12 | <.001 |
| Social cognition | 6.79 | .001 | -0.12 | 0.05 | -2.21 | .042 | 0.09 | 0.06 | 1.62 | .106 | -0.21 | 0.06 | -3.65 | .001 |
| Episodic memory | 7.21 | .001 | -0.12 | 0.06 | -1.93 | .054 | 0.13 | 0.06 | 2.03 | .054 | -0.25 | 0.07 | -3.79 | <.001 |
| Psychopathology factors^ | ***F*** | $p_{fdr}$ | ***B*** | ***SE*** | ***t*** | $p_{fdr}$ | ***B*** | ***SE*** | ***t*** | $p_{fdr}$ | ***B*** | ***SE*** | ***t*** | $p_{fdr}$ |
| Anxious-misery | 18.38 | <.001 | 0.31 | 0.07 | 4.70 | <.001 | 0.39 | 0.07 | 5.56 | <.001 | -0.08 | 0.07 | -1.15 | .249 |
| Psychosis | 12.17 | <.001 | 0.34 | 0.07 | 4.74 | <.001 | 0.25 | 0.08 | 3.38 | .001 | 0.08 | 0.08 | 1.05 | .292 |
| Behavioral | 25.46 | <.001 | 0.47 | 0.07 | 7.14 | <.001 | 0.23 | 0.07 | 3.27 | .001 | 0.24 | 0.07 | 3.38 | .001 |
| Fear | 39.66 | <.001 | 0.58 | 0.07 | 8.72 | <.001 | 0.39 | 0.07 | 5.59 | <.001 | 0.18 | 0.07 | 2.55 | .011 |
| Overall psychopathology | 250.30 | <.001 | 1.28 | 0.06 | 21.02 | <.001 | 1.07 | 0.06 | 16.45 | <.001 | 0.21 | 0.07 | 3.20 | .001 |

*df=1,138; ^df=1,136

**Supplementary Table 2. Group differences in volume and cortical thickness (n=1,141, df=1,134).**

| | | | S1-TD | | | | S2-TD | | | | S1-S2 | | | |
|---|---|---|---|---|---|---|---|---|---|---|---|---|---|---|
| | ***F*** | ***p*** | ***B*** | ***SE*** | ***t*** | $p_{fdr}$ | ***B*** | ***SE*** | ***t*** | $p_{fdr}$ | ***B*** | ***SE*** | ***t*** | $p_{fdr}$ |
| Total gray matter volume | 291.91 | <.001 | -49.75 | 3.23 | -15.42 | <.001 | 31.86 | 3.42 | 9.32 | <.001 | -81.61 | 3.44 | -23.70 | <.001 |
| Average cortical thickness | 151.43 | <.001 | -0.12 | 0.01 | -11.06 | <.001 | 0.08 | 0.01 | 6.79 | <.001 | -0.19 | .01 | -17.08 | <.001 |

#### Supplementary Table 3. Group differences in resting-state ALFF (n=840, df=834).

| | | | S1-TD | | | | S2-TD | | | | S1-S2 | | | |
|---|---|---|---|---|---|---|---|---|---|---|---|---|---|---|
| | *F* | *p* | *B* | *SE* | *t* | $p_{fdr}$ | *B* | *SE* | *t* | $p_{fdr}$ | *B* | *SE* | *t* | $p_{fdr}$ |
| Average resting-state ALFF | 17.09 | <.001 | -0.11 | 0.03 | -3.49 | .001 | 0.08 | 0.03 | 2.52 | .012 | -0.20 | 0.03 | -5.79 | <.001 |
| **Region** | *F* | $p_{fdr}$ | *B* | *SE* | *t* | $p_{fdr}$ | *B* | *SE* | *t* | $p_{fdr}$ | *B* | *SE* | *t* | $p_{fdr}$ |
| Right amygdala | 12.08 | <.001 | -0.15 | 0.04 | -4.12 | <.001 | 0.02 | 0.04 | 0.46 | .649 | -0.17 | 0.04 | -4.33 | <.001 |
| Left amygdala | 4.77 | .029 | -0.11 | 0.04 | -3.02 | .008 | -0.03 | 0.04 | -0.82 | .413 | -0.08 | 0.04 | -2.03 | .065 |
| Right hippocampus | 6.26 | .008 | -0.06 | 0.02 | -2.66 | .012 | 0.02 | 0.02 | 0.82 | .413 | -0.08 | 0.02 | -3.33 | .003 |
| Right anterior cingulate gyrus | 23.87 | <.001 | -0.25 | 0.05 | -5.52 | <.001 | 0.05 | 0.05 | 1.11 | .266 | -0.30 | 0.05 | -6.29 | <.001 |
| Left anterior cingulate gyrus | 19.22 | <.001 | -0.22 | 0.05 | -4.89 | <.001 | 0.05 | 0.05 | 1.10 | .270 | -0.28 | 0.05 | -5.69 | <.001 |
| Right anterior insula | 5.30 | .018 | -0.10 | 0.04 | -2.42 | .024 | 0.04 | 0.04 | 0.81 | .418 | -0.14 | 0.05 | -3.07 | .007 |
| Right anterior orbital gyrus | 12.52 | <.001 | -0.08 | 0.05 | -1.61 | .109 | 0.17 | 0.05 | 3.44 | .001 | -0.25 | 0.05 | -4.93 | <.001 |
| Left anterior orbital gyrus | 4.53 | .033 | -0.03 | 0.04 | -0.76 | .446 | 0.10 | 0.05 | 2.23 | .039 | -0.13 | 0.05 | -2.92 | .011 |
| Right angular gyrus | 4.73 | .029 | -0.09 | 0.05 | -1.73 | .126 | 0.08 | 0.05 | 1.45 | .147 | -0.16 | 0.05 | -3.06 | .007 |
| Left angular gyrus | 5.17 | .020 | -0.12 | 0.05 | -2.22 | .040 | 0.06 | 0.06 | 1.03 | .301 | -0.17 | 0.06 | -3.11 | .006 |
| Right entorhinal area | 4.29 | .040 | -0.10 | 0.04 | -2.68 | .023 | -0.01 | 0.04 | -0.21 | .836 | -0.09 | 0.04 | -2.34 | .030 |
| Right frontal operculum | 8.50 | .001 | -0.13 | 0.05 | -2.65 | .012 | 0.08 | 0.05 | 1.55 | .122 | -0.21 | 0.05 | -4.04 | <.001 |
| Right frontal pole | 12.46 | <.001 | -0.12 | 0.04 | -2.88 | .006 | 0.10 | 0.04 | 2.24 | .025 | -0.22 | 0.04 | -4.96 | <.001 |
| Left frontal pole | 12.22 | <.001 | -0.14 | 0.04 | -3.27 | .002 | 0.08 | 0.05 | 1.73 | .084 | -0.22 | 0.05 | -4.83 | <.001 |
| Right gyrus rectus | 6.49 | .007 | -0.06 | 0.03 | -1.69 | .092 | 0.07 | 0.03 | 2.04 | .063 | -0.13 | 0.03 | -3.60 | .001 |
| Left gyrus rectus | 5.98 | .010 | -0.05 | 0.04 | -1.44 | .149 | 0.08 | 0.04 | 2.10 | .054 | -0.13 | 0.04 | -3.45 | .002 |
| Right lateral orbital gyrus | 12.75 | <.001 | -0.03 | 0.04 | -0.64 | .522 | 0.18 | 0.04 | 4.12 | <.001 | -0.20 | 0.04 | -4.69 | <.001 |
| Left lateral orbital gyrus | 8.23 | .002 | -0.01 | 0.05 | -0.22 | .824 | 0.16 | 0.05 | 3.47 | .001 | -0.17 | 0.05 | -3.65 | .001 |
| Right medial frontal cortex | 14.69 | <.001 | -0.16 | 0.05 | -3.46 | .001 | 0.10 | 0.05 | 2.09 | .037 | -0.26 | 0.05 | -5.32 | <.001 |
| Left medial frontal cortex | 14.33 | <.001 | -0.14 | 0.05 | -2.99 | .004 | 0.13 | 0.05 | 2.53 | .011 | -0.27 | 0.05 | -5.33 | <.001 |
| Right middle frontal gyrus | 16.38 | <.001 | -0.20 | 0.06 | -3.51 | .001 | 0.14 | 0.06 | 2.34 | .019 | -0.34 | 0.06 | -5.66 | <.001 |
| Left middle frontal gyrus | 12.23 | <.001 | -0.18 | 0.06 | -3.07 | .003 | 0.12 | 0.06 | 1.98 | .048 | -0.30 | 0.06 | -4.88 | <.001 |
| Right medial orbital gyrus | 6.98 | .004 | -0.06 | 0.03 | -1.98 | .063 | 0.05 | 0.03 | 1.86 | .063 | -0.11 | 0.03 | -3.73 | .001 |
| Left medial orbital gyrus | 5.49 | .015 | -0.06 | 0.03 | -2.01 | .068 | 0.04 | 0.03 | 1.38 | .167 | -0.10 | 0.03 | -3.28 | .003 |
| Right superior frontal gyrus medial segment | 18.03 | <.001 | -0.26 | 0.06 | -4.58 | <.001 | 0.07 | 0.06 | 1.28 | .202 | -0.33 | 0.06 | -5.62 | <.001 |
| Left superior frontal gyrus medial segment | 15.85 | <.001 | -0.25 | 0.06 | -4.34 | <.001 | 0.07 | 0.06 | 1.12 | .262 | -0.32 | 0.06 | -5.24 | <.001 |
| Right middle temporal gyrus | 4.55 | .033 | -0.05 | 0.04 | -1.25 | .211 | 0.08 | 0.04 | 1.85 | .097 | -0.13 | 0.04 | -3.01 | .008 |
| Right opercular part of the inferior frontal gyrus | 12.83 | <.001 | -0.16 | 0.06 | -2.72 | .010 | 0.15 | 0.06 | 2.49 | .013 | -0.31 | 0.06 | -5.06 | <.001 |
| Left opercular part of the inferior frontal gyrus | 8.33 | .001 | -0.14 | 0.06 | -2.26 | .037 | 0.12 | 0.06 | 1.95 | .052 | -0.26 | 0.06 | -4.07 | <.001 |
| Right orbital part of the inferior frontal gyrus | 5.70 | .013 | -0.07 | 0.05 | -1.56 | .118 | 0.09 | 0.05 | 1.91 | .085 | -0.16 | 0.05 | -3.38 | .002 |
| Left orbital part of the inferior frontal gyrus | 4.51 | .033 | -0.05 | 0.05 | -0.91 | .365 | 0.12 | 0.05 | 2.10 | .054 | -0.16 | 0.06 | -2.94 | .010 |
| Right parahippocampal gyrus | 4.10 | .046 | -0.05 | 0.03 | -1.85 | .097 | 0.03 | 0.03 | 1.06 | .290 | -0.08 | 0.03 | -2.81 | .015 |
| Right posterior orbital gyrus | 6.21 | .008 | -0.07 | 0.03 | -2.38 | .026 | 0.04 | 0.03 | 1.20 | .229 | -0.11 | 0.03 | -3.42 | .002 |
| Right planum polare | 6.96 | .004 | -0.07 | 0.05 | -1.38 | .169 | 0.12 | 0.05 | 2.42 | .024 | -0.19 | 0.05 | -3.70 | .001 |
| Right superior frontal gyrus | 10.61 | <.001 | -0.17 | 0.05 | -3.17 | .002 | 0.08 | 0.06 | 1.46 | .146 | -0.25 | 0.06 | -4.46 | <.001 |
| Left superior frontal gyrus | 7.29 | .004 | -0.14 | 0.05 | -2.52 | .018 | 0.08 | 0.06 | 1.35 | .178 | -0.22 | 0.06 | -3.73 | .001 |
| Left superior parietal lobule | 4.15 | .045 | -0.10 | 0.05 | -1.79 | .111 | 0.07 | 0.06 | 1.17 | .243 | -0.16 | 0.06 | -2.84 | .014 |
| Right temporal pole | 15.16 | <.001 | -0.11 | 0.03 | -3.97 | <.001 | 0.05 | 0.03 | 1.54 | .124 | -0.16 | 0.03 | -5.25 | <.001 |
| Left temporal pole | 9.58 | <.001 | -0.10 | 0.03 | -3.22 | .002 | 0.03 | 0.03 | 1.11 | .268 | -0.13 | 0.03 | -4.15 | <.001 |
| Right triangular part of the inferior frontal gyrus | 16.75 | <.001 | -0.15 | 0.05 | -2.87 | .004 | 0.17 | 0.06 | 3.08 | .003 | -0.32 | 0.06 | -5.79 | <.001 |
| Left triangular part of the inferior frontal gyrus | 7.50 | .003 | -0.09 | 0.06 | -1.63 | .104 | 0.14 | 0.06 | 2.34 | .029 | -0.24 | 0.06 | -3.86 | <.001 |

**Supplementary Table 4. Group differences in fractional anisotropy of white matter tracts (n=923, df=916).**

| | | | S1-TD | | | | S2-TD | | | | S1-S2 | | | |
|---|---|---|---|---|---|---|---|---|---|---|---|---|---|---|
| | ***F*** | ***p*** | ***B*** | ***SE*** | ***t*** | $p_{fdr}$ | ***B*** | ***SE*** | ***t*** | $p_{fdr}$ | ***B*** | ***SE*** | ***t*** | $p_{fdr}$ |
| Average fractional anisotropy | 18.65 | <.001 | -0.21 | 0.05 | -4.63 | <.001 | 0.06 | 0.05 | 1.34 | .179 | -0.28 | 0.05 | -5.71 | <.001 |
| **Tract** | ***F*** | $p_{fdr}$ | ***B*** | ***SE*** | ***t*** | $p_{fdr}$ | ***B*** | ***SE*** | ***t*** | $p_{fdr}$ | ***B*** | ***SE*** | ***t*** | $p_{fdr}$ |
| Left anterior thalamic radiation | 7.09 | .002 | -0.08 | 0.06 | -1.23 | .217 | 0.17 | 0.07 | 2.56 | .016 | -0.25 | 0.07 | -3.72 | .001 |
| Right anterior thalamic radiation | 5.74 | .007 | -0.08 | 0.06 | -1.29 | .198 | 0.15 | 0.07 | 2.16 | .047 | -0.23 | 0.07 | -3.37 | .002 |
| Right corticospinal tract | 3.61 | .049 | -0.15 | 0.07 | -2.21 | .041 | 0.02 | 0.07 | 0.32 | .751 | -0.18 | 0.07 | -2.40 | .041 |
| Left cingulum (hippocampus) | 9.88 | <.001 | -0.24 | 0.07 | -3.44 | .001 | 0.07 | 0.07 | 0.89 | .373 | -0.31 | 0.07 | -4.11 | <.001 |
| Right cingulum (hippocampus) | 8.66 | .001 | -0.22 | 0.07 | -3.01 | .004 | 0.09 | 0.08 | 1.18 | .240 | -0.31 | 0.08 | -3.96 | <.001 |
| Forceps minor | 10.70 | <.001 | -0.22 | 0.07 | -3.30 | .002 | 0.09 | 0.07 | 1.32 | .187 | -0.31 | 0.07 | -4.42 | <.001 |
| Left inferior longitudinal fasciculus | 10.66 | <.001 | -0.29 | 0.07 | -3.96 | <.001 | 0.02 | 0.08 | 0.24 | .812 | -0.31 | 0.08 | -3.97 | <.001 |
| Right inferior longitudinal fasciculus | 18.82 | <.001 | -0.34 | 0.07 | -4.60 | <.001 | 0.11 | 0.08 | 1.42 | .155 | -0.46 | 0.08 | -5.76 | <.001 |
| Right superior longitudinal fasciculus | 7.13 | .002 | -0.23 | 0.07 | -3.52 | .001 | -0.03 | 0.07 | -0.46 | .644 | -0.20 | 0.07 | -2.87 | .006 |
| Left uncinate fasciculus | 5.96 | .006 | -0.23 | 0.07 | -3.16 | .005 | -0.02 | 0.08 | -0.24 | .811 | -0.21 | 0.08 | -2.73 | .010 |

**Supplementary Table 5. Demographics, cognition, academic achievement, and psychopathology sensitivity analyses that exclude those taking psychotropic medications (n=1,037).**

| | | | S1-TD | | | | S2-TD | | | | S1-S2 | | | |
|---|---|---|---|---|---|---|---|---|---|---|---|---|---|---|
| | $F/X^2$ | *p* | *B* | *SE* | *t* | $p_{fdr}$ | *B* | *SE* | *t* | $p_{fdr}$ | *B* | *SE* | *t* | $p_{fdr}$ |
| Age* | 4.36 | .013 | 0.79 | 0.27 | 2.95 | .010 | 0.35 | 0.29 | 1.22 | .222 | 0.44 | 0.30 | 1.47 | .212 |
| Gender* | 15.10 | <.001 | 0.55 | 0.15 | 3.72 | .001 | 0.40 | 0.16 | 2.54 | .017 | 0.15 | 0.16 | 0.90 | .369 |
| Maternal level of education* | 16.90 | <.001 | -0.95 | 0.17 | -5.45 | <.001 | -0.10 | 0.19 | -0.53 | .595 | -0.85 | 0.19 | -4.39 | <.001 |
| Academic achievement^ | 30.27 | <.001 | -7.80 | 1.16 | -6.74 | <.001 | 0.69 | 1.24 | 0.56 | .579 | -8.49 | 1.27 | -6.66 | <.001 |
| Overall accuracy^ | 22.54 | <.001 | -0.27 | 0.05 | -5.13 | <.001 | 0.09 | 0.06 | 1.65 | .098 | -0.37 | 0.06 | -6.27 | <.001 |
| Accuracy factors^ | *F* | $p_{fdr}$ | *B* | *SE* | *t* | $p_{fdr}$ | *B* | *SE* | *t* | $p_{fdr}$ | *B* | *SE* | *t* | $p_{fdr}$ |
| Executive function/complex reasoning | 32.74 | <.001 | -0.37 | 0.06 | -6.56 | <.001 | 0.08 | 0.06 | 1.39 | .163 | -0.45 | 0.06 | -7.32 | <.001 |
| Social cognition | 3.55 | .029 | -0.08 | 0.06 | -1.39 | .165 | 0.09 | 0.06 | 1.44 | .165 | -0.16 | 0.06 | -2.66 | .024 |
| Episodic memory | 7.88 | .001 | -0.13 | 0.06 | -1.99 | .047 | 0.15 | 0.07 | 2.22 | .040 | -0.28 | 0.07 | -3.97 | <.001 |
| Psychopathology factors^ | *F* | $p_{fdr}$ | *B* | *SE* | *t* | $p_{fdr}$ | *B* | *SE* | *t* | $p_{fdr}$ | *B* | *SE* | *t* | $p_{fdr}$ |
| Anxious-misery | 15.58 | <.001 | 0.30 | 0.07 | 4.56 | <.001 | 0.35 | 0.07 | 4.89 | <.001 | -0.05 | 0.07 | -0.62 | .533 |
| Psychosis | 12.08 | <.001 | 0.34 | 0.07 | 4.59 | <.001 | 0.28 | 0.08 | 3.58 | .001 | 0.06 | 0.08 | 0.68 | .496 |
| Behavioral | 21.46 | <.001 | 0.42 | 0.07 | 6.47 | <.001 | 0.13 | 0.07 | 1.90 | .057 | 0.29 | 0.07 | 4.02 | <.001 |
| Fear | 41.80 | <.001 | 0.61 | 0.07 | 8.88 | <.001 | 0.43 | 0.07 | 5.83 | <.001 | 0.18 | 0.08 | 2.39 | .017 |
| Overall psychopathology | 218.31 | <.001 | 1.24 | 0.06 | 19.54 | <.001 | 1.04 | 0.07 | 15.20 | <.001 | 0.21 | 0.07 | 2.95 | .003 |

*df=1,034; ^df=1,032

**Supplementary Table 6. Volume and cortical thickness sensitivity analyses that exclude those taking psychotropic medications (n=1,037, df=1,030).**

| | | | S1-TD | | | | S2-TD | | | | S1-S2 | | | |
|---|---|---|---|---|---|---|---|---|---|---|---|---|---|---|
| | ***F*** | ***p*** | ***B*** | ***SE*** | ***t*** | $p_{fdr}$ | ***B*** | ***SE*** | ***t*** | $p_{fdr}$ | ***B*** | ***SE*** | ***t*** | $p_{fdr}$ |
| Total gray matter volume | 249.86 | <.001 | -48.23 | 3.38 | -14.27 | <.001 | 32.75 | 3.59 | 9.12 | <.001 | -80.98 | 3.69 | -21.97 | <.001 |
| Average cortical thickness | 141.27 | <.001 | -0.12 | 0.01 | -10.99 | <.001 | 0.08 | 0.01 | 6.55 | <.001 | -0.20 | 0.01 | -16.45 | <.001 |

**Supplementary Table 7. Resting-state ALFF sensitivity analyses that exclude those taking psychotropic medications (n=763, df=756).**

| | | | S1-TD | | | | S2-TD | | | | S1-S2 | | | |
|---|---|---|---|---|---|---|---|---|---|---|---|---|---|---|
| | *F* | *p* | *B* | *SE* | *t* | $p_{fdr}$ | *B* | *SE* | *t* | $p_{fdr}$ | *B* | *SE* | *t* | $p_{fdr}$ |
| Average resting-state ALFF | 13.90 | <.001 | -0.12 | 0.04 | -3.26 | .001 | 0.09 | 0.04 | 2.25 | .025 | -0.21 | 0.04 | -5.21 | <.001 |
| **Region** | *F* | $p_{fdr}$ | *B* | *SE* | *t* | $p_{fdr}$ | *B* | *SE* | *t* | $p_{fdr}$ | *B* | *SE* | *t* | $p_{fdr}$ |
| Right amygdala | 8.89 | .001 | -0.14 | 0.04 | -3.67 | .001 | 0.01 | 0.04 | 0.23 | .818 | -0.15 | 0.04 | -3.60 | .001 |
| Left amygdala | 4.53 | .046 | -0.12 | 0.04 | -2.95 | .010 | -0.03 | 0.04 | -0.78 | .437 | -0.08 | 0.04 | -1.94 | .079 |
| Right anterior cingulate gyrus | 19.44 | <.001 | -0.25 | 0.05 | -5.31 | <.001 | 0.03 | 0.05 | 0.57 | .568 | -0.28 | 0.05 | -5.42 | <.001 |
| Left anterior cingulate gyrus | 14.40 | <.001 | -0.21 | 0.05 | -4.51 | <.001 | 0.03 | 0.05 | 0.61 | .545 | -0.24 | 0.05 | -4.72 | <.001 |
| Right anterior orbital gyrus | 11.48 | <.001 | -0.07 | 0.05 | -1.48 | .140 | 0.18 | 0.05 | 3.43 | .001 | -0.26 | 0.05 | -4.70 | <.001 |
| Right frontal operculum | 6.54 | .009 | -0.12 | 0.05 | -2.30 | .033 | 0.08 | 0.06 | 1.47 | .142 | -0.20 | 0.06 | -3.56 | .001 |
| Right frontal pole | 9.18 | .001 | -0.11 | 0.04 | -2.54 | .017 | 0.09 | 0.05 | 1.95 | .051 | -0.20 | 0.05 | -4.25 | <.001 |
| Left frontal pole | 8.48 | .002 | -0.13 | 0.05 | -2.73 | .010 | 0.07 | 0.05 | 1.53 | .127 | -0.20 | 0.05 | -4.02 | <.001 |
| Right gyrus rectus | 4.65 | .042 | -0.04 | 0.03 | -1.22 | .223 | 0.07 | 0.04 | 1.97 | .075 | -0.11 | 0.04 | -3.04 | .007 |
| Right lateral orbital gyrus | 12.15 | <.001 | -0.02 | 0.04 | -0.51 | .608 | 0.19 | 0.05 | 4.15 | <.001 | -0.22 | 0.05 | -4.53 | <.001 |
| Left lateral orbital gyrus | 7.13 | .005 | 0.00 | 0.05 | -0.02 | .987 | 0.17 | 0.05 | 3.37 | .001 | -0.17 | 0.05 | -3.30 | .001 |
| Right medial frontal cortex | 10.30 | <.001 | -0.15 | 0.05 | -3.14 | .003 | 0.08 | 0.05 | 1.55 | .122 | -0.23 | 0.05 | -4.39 | <.001 |
| Left medial frontal cortex | 10.30 | <.001 | -0.12 | 0.05 | -2.57 | .016 | 0.11 | 0.05 | 2.21 | .027 | -0.24 | 0.05 | -4.52 | <.001 |
| Right middle frontal gyrus | 10.48 | <.001 | -0.16 | 0.06 | -2.80 | .008 | 0.12 | 0.06 | 1.99 | .047 | -0.29 | 0.06 | -4.53 | <.001 |
| Left middle frontal gyrus | 7.86 | .003 | -0.15 | 0.06 | -2.40 | .025 | 0.11 | 0.06 | 1.75 | .081 | -0.26 | 0.07 | -3.93 | <.001 |
| Right medial orbital gyrus | 5.43 | .024 | -0.05 | 0.03 | -1.68 | .094 | 0.06 | 0.03 | 1.78 | .094 | -0.11 | 0.03 | -3.29 | .003 |
| Right superior frontal gyrus medial segment | 13.14 | <.001 | -0.24 | 0.06 | -4.11 | <.001 | 0.06 | 0.06 | 0.90 | .369 | -0.30 | 0.06 | -4.68 | <.001 |
| Left superior frontal gyrus medial segment | 10.79 | <.001 | -0.23 | 0.06 | -3.82 | <.001 | 0.04 | 0.06 | 0.65 | .516 | -0.27 | 0.06 | -4.17 | <.001 |
| Right opercular part of the inferior frontal gyrus | 9.27 | .001 | -0.14 | 0.06 | -2.24 | .026 | 0.15 | 0.06 | 2.29 | .026 | -0.28 | 0.07 | -4.30 | <.001 |
| Left opercular part of the inferior frontal gyrus | 5.11 | .030 | -0.10 | 0.06 | -1.65 | .099 | 0.11 | 0.07 | 1.71 | .099 | -0.22 | 0.07 | -3.19 | .004 |
| Right orbital part of the inferior frontal gyrus | 4.70 | .042 | -0.06 | 0.05 | -1.17 | .244 | 0.10 | 0.05 | 2.01 | .067 | -0.16 | 0.05 | -3.04 | .007 |
| Right posterior orbital gyrus | 5.26 | .027 | -0.07 | 0.03 | -2.28 | .034 | 0.04 | 0.03 | 1.06 | .290 | -0.11 | 0.04 | -3.12 | .006 |
| Right superior frontal gyrus | 6.60 | .009 | -0.15 | 0.06 | -2.63 | .013 | 0.06 | 0.06 | 1.06 | .291 | -0.21 | 0.06 | -3.47 | .002 |
| Right temporal pole | 11.86 | <.001 | -0.11 | 0.03 | -3.63 | <.001 | 0.04 | 0.03 | 1.30 | .193 | -0.15 | 0.03 | -4.60 | <.001 |
| Left temporal pole | 7.54 | .004 | -0.09 | 0.03 | -2.83 | .007 | 0.04 | 0.03 | 1.10 | .270 | -0.13 | 0.03 | -3.70 | .001 |
| Right triangular part of the inferior frontal gyrus | 11.59 | <.001 | -0.13 | 0.06 | -2.37 | .018 | 0.16 | 0.06 | 2.68 | .011 | -0.29 | 0.06 | -4.82 | <.001 |
| Left triangular part of the inferior frontal gyrus | 4.98 | .033 | -0.06 | 0.06 | -1.00 | .316 | 0.14 | 0.06 | 2.23 | .039 | -0.21 | 0.07 | -3.10 | .006 |

**Supplementary Table 8. Fractional anisotropy sensitivity analyses that exclude those taking psychotropic medications (n=836, df=828).**

| | | | S1-TD | | | | S2-TD | | | | S1-S2 | | | |
|---|---|---|---|---|---|---|---|---|---|---|---|---|---|---|
| | ***F*** | ***p*** | ***B*** | ***SE*** | ***t*** | $p_{fdr}$ | ***B*** | ***SE*** | ***t*** | $p_{fdr}$ | ***B*** | ***SE*** | ***t*** | $p_{fdr}$ |
| Average fractional anisotropy | 17.17 | <.001 | -0.23 | 0.05 | -4.62 | <.001 | 0.06 | 0.05 | 1.15 | .249 | -0.29 | 0.05 | -5.39 | <.001 |
| **Tract** | ***F*** | $p_{fdr}$ | ***B*** | ***SE*** | ***t*** | $p_{fdr}$ | ***B*** | ***SE*** | ***t*** | $p_{fdr}$ | ***B*** | ***SE*** | ***t*** | $p_{fdr}$ |
| Left anterior thalamic radiation | 5.17 | .012 | -0.07 | 0.07 | -1.03 | .301 | 0.16 | 0.07 | 2.25 | .037 | -0.23 | 0.07 | -3.17 | .005 |
| Right anterior thalamic radiation | 5.42 | .010 | -0.09 | 0.07 | -1.30 | .192 | 0.16 | 0.07 | 2.11 | .052 | -0.25 | 0.07 | -3.28 | .003 |
| Left cingulum (hippocampus) | 8.16 | .001 | -0.23 | 0.07 | -3.14 | .003 | 0.07 | 0.08 | 0.88 | .381 | -0.30 | 0.08 | -3.74 | .001 |
| Right cingulum (hippocampus) | 8.59 | .001 | -0.25 | 0.08 | -3.29 | .002 | 0.07 | 0.08 | 0.82 | .414 | -0.32 | 0.08 | -3.80 | <.001 |
| Forceps minor | 9.59 | <.001 | -0.22 | 0.07 | -3.16 | .002 | 0.09 | 0.07 | 1.29 | .197 | -0.31 | 0.07 | -4.18 | <.001 |
| Left inferior longitudinal fasciculus | 10.25 | <.001 | -0.31 | 0.08 | -3.97 | <.001 | 0.01 | 0.08 | 0.17 | .867 | -0.32 | 0.08 | -3.82 | <.001 |
| Right inferior longitudinal fasciculus | 17.84 | <.001 | -0.36 | 0.08 | -4.61 | <.001 | 0.11 | 0.08 | 1.33 | .184 | -0.47 | 0.09 | -5.55 | <.001 |
| Right superior longitudinal fasciculus | 7.41 | .002 | -0.25 | 0.07 | -3.64 | .001 | -0.04 | 0.07 | -0.51 | .607 | -0.21 | 0.08 | -2.85 | .007 |
| Left uncinate fasciculus | 5.77 | .008 | -0.25 | 0.08 | -3.32 | .003 | -0.07 | 0.08 | -0.83 | .406 | -0.18 | 0.08 | -2.24 | .038 |

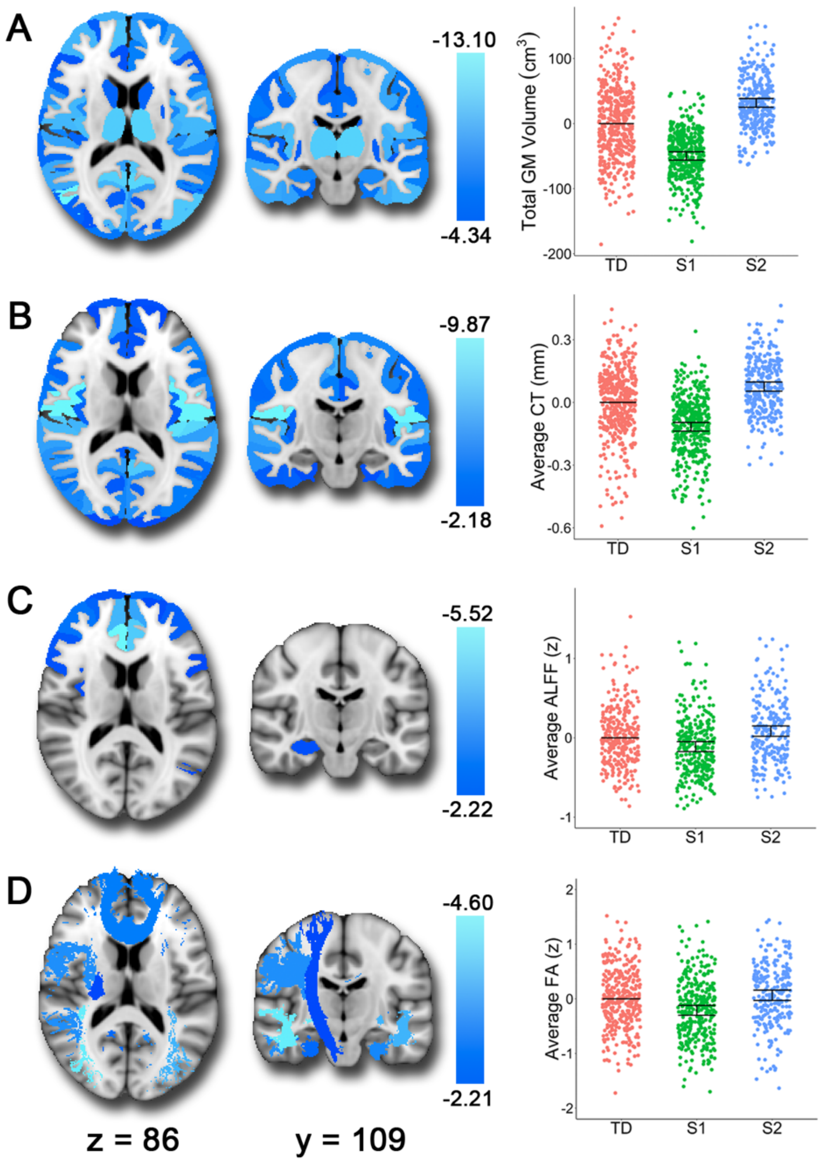

**Supplementary Figure 1. Subtype 1 (S1) shows smaller volume, thinner cortex, lower resting-state ALFF, and reduced white matter integrity relative to typically developing youth (TD).** The brain images show the t-values for the S1>TD contrast. In the scatterplots, we show the estimates from the fitted GAM model with all three groups for comparison. Each vertical line represents the 95% confidence interval (CI), with the comparison group (TD) represented by its mean line. The subtype is significantly different from TD if its corresponding CI does not contain 0 (the mean of TD). In comparison to TD, S1 showed **A)** smaller volumes, **B)** reduced cortical thickness, **C)** reduced resting-state ALFF (amplitude of low frequency fluctuations) in frontal regions, bilateral amygdala, and right hippocampus, and **D)** reduced fractional anisotropy in white matter tracts including the inferior longitudinal fasciculi, uncinate fasciculus, corticospinal tract, parahippocampal cingulum, superior longitudinal fasciculus, and forceps minor.

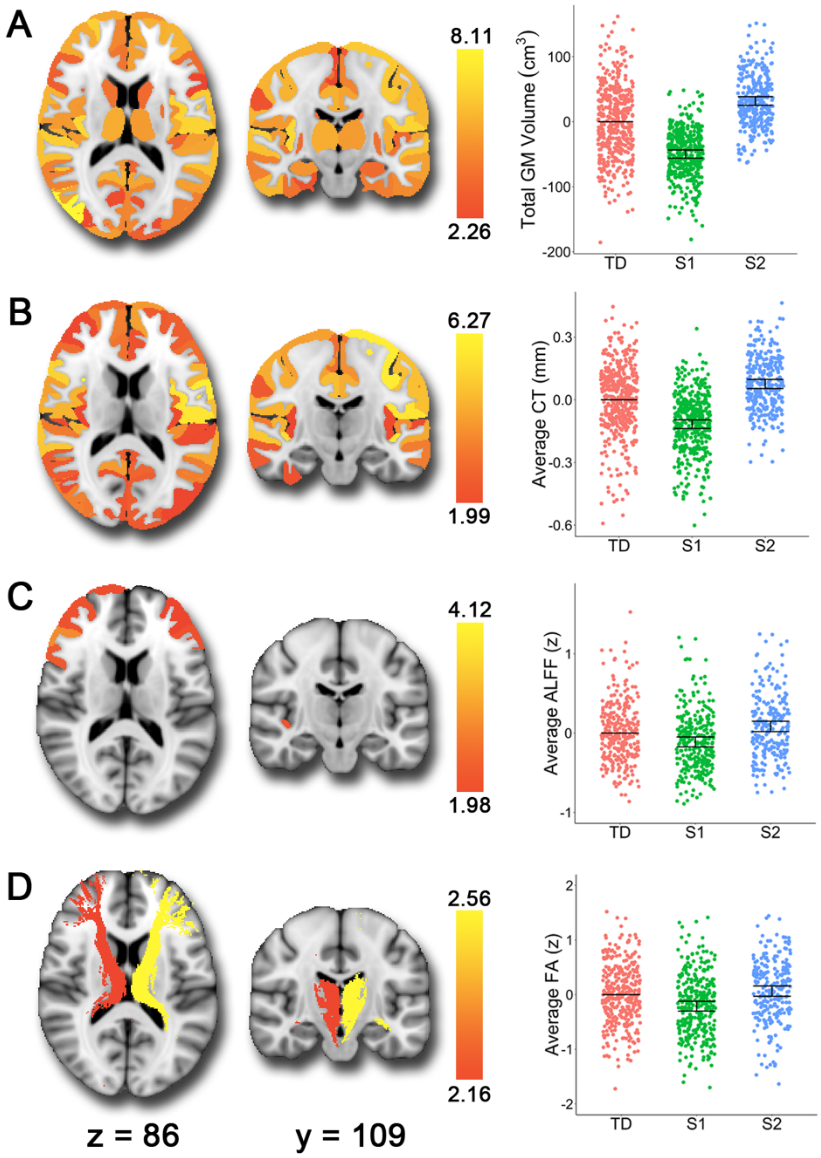

**Supplementary Figure 2. Subtype 2 (S2) shows larger volume, thicker cortex, higher resting-state ALFF, and greater white matter integrity relative to typically developing youth (TD).** The brain images show the t-values for the S2>TD contrast. In the scatterplots, we show the estimates from the fitted GAM model with all three groups for comparison. Each vertical line represents the 95% confidence interval (CI), with the comparison group (TD) represented by its mean line. The subtype is significantly different from TD if its corresponding CI does not contain 0 (the mean of TD). In comparison to TD, S2 showed **A)** larger volumes, **B)** greater cortical thickness, **C)** higher resting-state ALFF (amplitude of low frequency fluctuations) in frontal regions, and **D)** greater fractional anisotropy in the left and right anterior thalamic radiations.
